# To be and not to be: Wide-field Ca^2+^ imaging reveals neocortical functional segmentation combines stability and flexibility

**DOI:** 10.1101/2022.09.16.508301

**Authors:** Angela K. Nietz, Martha L. Streng, Laurentiu S. Popa, Russell E. Carter, Evelyn Flaherty, Justin D. Aronson, Timothy J. Ebner

## Abstract

The stability and flexibility of the functional parcellation of the cerebral cortex is fundamental to how familiar and novel information is both represented and stored. We leveraged new advances in Ca^2+^ sensors and microscopy to understand the dynamics of functional segmentation in the dorsal cerebral cortex. We performed wide-field Ca^2+^ imaging in head-fixed mice and used spatial Independent Component Analysis (ICA) to identify independent spatial sources of Ca^2+^ fluorescence. The imaging data were evaluated over multiple timescales and discrete behaviors including resting, walking, and grooming. When evaluated over the entire dataset, a set of template independent components (ICs) were identified that were common across behaviors. Template ICs were present across a range of timescales, from days to 30 seconds, although with lower occurrence probability at shorter timescales, highlighting the stability of the functional segmentation. Importantly, unique ICs emerged at the shorter duration timescales that could act to transiently refine the cortical network. When data were evaluated by behavior, both common and behavior-specific ICs emerged. Each behavior is composed of unique combinations of common and behavior-specific ICs. These observations suggest that cerebral cortical functional segmentation exhibits considerable spatial stability over time and behaviors while retaining the flexibility for task-dependent reorganization.

## Introduction

Animal behavior is rich, dynamic, and expressed in a high dimensional space. Encoding of such a behavioral repertoire in the brain requires equally complex and flexible computations at multiple levels of processing. One of the prominent hypotheses regarding the nervous system is that behavior is an emerging property of neuronal populations across the brain (Buzsáki 2010; Yuste 2015; Makino et al. 2017). In this view, behaviorally relevant computations are distributed within and across anatomically and functionally distinct brain regions, at the level of both cells and circuits. Therefore, a comprehensive characterization of neuronal activity at various spatial and temporal scales, including simultaneously recording from different regions, is critical for deciphering how the brain plans, learns, and executes behaviors (Fox et al. 2005; Allen et al. 2017; vena-Koenigsberger et al. 2017; Gilad et al. 2018; Ren and Komiyama 2021).

Observing neuronal activity at the mesoscale level is fundamental to investigations into both the representation and control of behavior, as it allows simultaneous characterization of local neuronal population activity as well as long-range interactions among these local populations. The development of fast, genetically encoded calcium (Ca^2+^) sensors was critical for mesoscale imaging, as it allows simultaneous visualization of neural activity across brain regions at high spatiotemporal resolution and provides for cell-type specificity (Che and De Marco García 2021; Guo et al. 2021). When combined with newer techniques for large-scale, chronic cranial windows, which can give chronic access to 100 mm^2^ of cortex, wide-field Ca^2+^ imaging of neuronal activity is proving to be a powerful tool to investigate local and long-range ensemble interactions (Ghanbari et al. 2019; Cardin et al. 2020; Cramer et al. 2021; Ren and Komiyama 2021; Takahashi et al. 2021).

Wide-field Ca^2+^ imaging in the cerebral cortex has already provided insights into the spatial and temporal patterns of activation during behaviors, including reorganization of cortical interactions with learning (Makino et al. 2017; Gilad et al. 2018), contribution of spontaneous movements to cortical activity (Musall et al. 2019), state transitions and changes in functional connectivity during locomotion (West et al. 2021), cortical dynamics during reaching (Galiñanes et al. 2018), and how task information across the cortex is modulated by cognitive processes and task demands (Pinto et al. 2019; Salkoff et al. 2020; Zatka-Haas et al. 2021).

Given the recent prominence of Ca^2+^ imaging for investigating mesoscopic neuronal dynamics and functional interactions among ensembles and their interactions with behavior, many important questions focus on how the underlying activity is functionally parcellated in the cerebral cortex (Cardin et al. 2020). This makes it necessary to identify regions engaged in neural processing and then determine how they vary with time and behavior. Robust segmentation is critical to understand the regions engaged in behaviors and their interactions. One approach is to use existing anatomical and/or functional maps. Both seed-based correlational and Local Semi-nonnegative Matrix Factorization (LocaNMF) studies have utilized the mouse Allen Brain Atlas Common Cortical Framework (CCF) to segment the cerebral cortex (Barson et al. 2020; Saxena et al. 2020; Wang et al. 2020; Xiao et al. 2021; Quarta et al. 2022). While having many advantages, the use of static maps does not take into account individual variability, nor that segmentation may change with different behaviors and over time.

Another approach is blind-source separation with spatial Independent Component Analysis (ICA) (McKeown et al. 1998; Calhoun and Adali 2006). Spatial ICA separates a multivariate signal into additive subcomponents by maximizing statistical independence from each other, assuming that the subcomponents are non-Gaussian signals. ICA can be used to identify both temporal and spatial independent components (ICs). Extensively used in human brain imaging, spatial ICA has numerous advantages, including being data-based with minimal prior assumptions, high reliability, artifact detection/rejection, and repeatability across studies and subjects (Calhoun and de Lacy 2017).

To date, a few wide-field Ca^2+^ imaging studies have used ICA to segment the dorsal cerebral cortex into functional domains (Reidl et al. 2007; Makino et al. 2017; Weiser et al. 2021; West et al. 2021). In most cases, ICA was performed across the entire dataset, generating a singular set of ICs. It remains to be determined the degree to which ICs are stable across timescales and behaviors, or the degree to which they emerge and dissipate transiently in a behavior-dependent manner.

To address these questions, we examined several properties of spatial ICs obtained from wide-field Ca^2+^ imaging of excitatory cortical neurons in awake, behaving mice. First, we examine the similarity and repeatability of components obtained at different timescales. The results show that the ICs extracted at different timescales are spatially stable, although the probability of occurrence decreases at shorter duration time timescales. We also show that at shorter durations, unique ICs are found but at a relatively low probability. We also examine whether the ICs extracted depend on the behavior, comparing rest, locomotion, and grooming. The results demonstrate both a common set of ICs across these three behaviors as well as sets of unique ICs that arise during specific behaviors. These data highlight the need for both a subject-based analysis approach, to capture intra and inter-subject variability, and a task-based analysis approach, to capture changing neural dynamics.

## Materials and Methods

All experiments were approved by the Institutional Animal Care and Use Committee (IACUC) at the University of Minnesota.

### Animals, implant fabrication, and surgical procedures

Six mice of both sexes (ages 6-12 months) expressing the genetically encoded Ca^2+^ indicator GCaMP6f primarily in excitatory neurons (Thy1-GCaMP6f; Jackson Laboratories #024339) were used for the imaging studies (Dana et al. 2014). Animals were implanted with “See-Shell” transparent polymer skulls, designed to conform to the geometry of the skull and provide chronic optical access to a large contiguous region of the dorsal cerebral cortex, as previously detailed (Ghanbari et al. 2019; West et al. 2021). These polymer windows consist of 50 μm thick, transparent polyethylene terephthalate (PET) film (MELINEX 462, Dupont Inc.), fitted and bonded (Scotch-Weld™ DP100 Plus Clear, 3M Inc.) to a 3D-printed frame made from polymethylmethacrylate (PMMA) (RSF2-GPBK-04, Formlabs Inc.).

Implantation of the See-Shells was performed as described previously, including anesthesia, analgesia, monitoring, craniotomy, and post-operative care (Ghanbari et al. 2019; West et al. 2021). Following removal of the bone flap, the See-Shells were aligned over the brain at the edges of the craniotomy and attached to the skull with cyanoacrylate glue (Vetbond, 3M). See-Shells were further secured using dental cement (S380, C&B Metabond, Parkell Inc.) and a custom titanium head plate attached to the implant via three screws and also secured with dental cement.

### Experimental Setup

Following surgery, mice were housed singly on a 12-hour reverse light-dark cycle with experiments performed during the dark phase. Imaging experiments were performed with mice head-fixed to a freely moving disk treadmill that allows for a variety of behaviors including rest, walking, and grooming. Behavior was monitored and recorded using a high-speed IR-sensitive CMOS camera (40 fps; Blackfly, FLIR Systems) under infrared illumination that did not affect concurrent Ca^2+^ imaging. Spontaneous locomotion was also monitored using a rotary encoder attached to the disk treadmill. Wheel rotation was recorded at 1kHz using an Arduino microcontroller (Arduino Mega 2560; Arduino). Rotary encoder data was subsequently used to segment Ca^2+^ imaging data into periods of walk and rest, as previously described (West et al. 2021). In order to segment walk and rest data, we defined rest empirically as any movement less than 0.15 cm/s in either the forward or reverse direction and walk as any movement greater than 0.25 cm/s in the forward direction only (West et al. 2021).

In a subset of experiments, a waterspout was placed above the animal’s snout and a droplets of water delivered to encourage grooming behavior. Delivery of water droplets was controlled using a Bpod Finite State Machine (Bpod, Sanworks) and provided software as well as custom written Matlab code (Mathworks; Natick, MA). Grooming behavior was manually segmented from high-speed videos with a grooming bout manually defined as when the mouse first lifts its paw toward its snout and ending when the paw returns toward the wheel or is held stationary in an upright position longer than 1.5-2 seconds. Any grooming bouts split by less than ~1.5-2 seconds of walk or rest were combined into a single grooming period. Manually segmented grooming bouts were removed from walk and rest data using custom written Matlab code.

### Mesoscale Calcium Imaging

Cortex-wide mesoscale Ca^2+^ imaging was performed by placing head-fixed mice beneath an epifluorescence microscope (Nikon AZ-100) and focusing on layers II/III of the cerebral cortex. Dual-wavelength single photon imaging was performed using a fast LED switcher (OptoLED, Cairn). Calcium-dependent and independent signals of GCaMP6f were sampled by alternating illumination of a 470 nm and 405 nm LED, respectively. Mesoscale brain images were captured (40 fps; 18 ms exposure; 256 x 256 pixels; ~24 x 24 μm pixel size) using a high-speed CMOS camera (Andor Zyla, 4.2, Oxford Instruments) controlled by Micromanager imaging software (Edelstein et al. 2010; Edelstein et al. 2014). Using digital zoom, the field-of-view was adjusted so that See-Shell cortical window (~6.2 x 6.2 mm) filled as much of the field as possible. Continuous images were collected as tiff stacks over either 5.5- or 6-minute periods which comprised a single imaging trial, with each imaging day comprised of several trials (3-10 trials per mouse per day). Synchronization of behavior and microscope cameras as well as the rotary encoder was controlled by a series of TTL pulses delivered by Spike2 software in conjunction with a CED Power 1401 data acquisition system (Cambridge Electronic Design).

### Pre-processing of mesoscopic imaging data

For each individual trial, the first preprocessing step was to de-interleave into 470 nm and 405 nm channels and remove the initial 30 seconds of each trial to eliminate any initial rundown of the Ca^2+^ signal. All subsequent analyses were done on the remaining 5 or 5.5 minutes of data in a trial. Calcium-dependent GCaMP6f images (470 nm) were corrected for Ca^2+^-independent signals (405 nm images) using the following equation implemented as part of a custom Matlab code, similar to corrections described by others (Vanni and Murphy 2014; Jacobs et al. 2020; MacDowell and Buschman 2020; West et al. 2021).

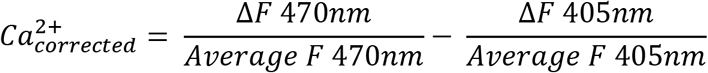

The hemodynamic corrected frames were saved along with the reference background 470 nm frame. Subsequently, a reference frame was chosen, and a mask was drawn for each animal to separate the brain from the non-brain field-of-view (FOV). For each mouse, all hemodynamic corrected trials were co-registered to the reference frame using a rigid transformation (*imregconfig* and *imregtform* functions in Matlab). For each mouse, the hemodynamic corrected frames were smoothed in the spatial domain using a 5 x 5 pixel Gaussian filter and concatenated across all recording days. The noise present in the brain FOV due to variability in the background illumination was eliminated at pixel level by regressing the data against the average of the non-brain pixels and performing all analyses on the residuals. The de-noised brain data were compressed using singular value decomposition (SVD), retaining the first 200 components (Musall et al. 2019; Saxena et al. 2020). Data for each animal were sub-sectioned at four different timescales: 1) days (15-50 minutes per day), 2) trials (5-5.5 minutes), 3) minutes, and 4) 30 seconds. At each timescale, concatenation and compression were repeated as described for the full dataset.

### Spatial independent component analysis

Spatial ICA was performed on the entire concatenated data set and on each of the sub-sectioned data sets, independently. For each of these data sets, we computed 60 independent components (ICs) using the Joint Approximation and Diagonalization of Eigenmatrices (JADE) algorithm that obtains maximally independent source signals from signal mixtures by minimizing mutual information (Cardoso 1999; Sahonero-Alvarez and Calderon 2017) as has been used in previous wide-field Ca^2+^ imaging studies (Makino et al. 2017; West et al. 2021). The solutions were multiplied back into the original vector space and their z-scores computed to yield spatial maps of the ICs. Binary masks of the ICs were obtained by setting values less than a z-score of 2.5 to 0 and all other values to 1. Spatial ICs with less than 250 contiguous pixels were excluded from analysis. Any IC containing more than one non-contiguous region, for example a pair of homotopic regions, was separated into the individual regions, so each IC mask consisted of a single, contiguous region. Remaining ICs were inspected for artifacts and were manually discarded, including areas overlying only vasculature (Musall et al. 2019; West et al. 2021).

### Comparison of ICA segmentation across timescales

The first analysis focused on determining matches among the ICs obtained at different timescales. For each mouse, the catalog of ICs obtained from the entire concatenated dataset served as the ground-truth template map to which all other ICs obtained at the different timescales were compared. ICs were sorted anterior to posterior based on the IC centroid position and assigned a unique color value. Pairwise comparisons of ICs in the template map and timescale maps were performed using the Jaccard index, a common and robust statistic to determine similarity between data sets, including images and more recently graphs and segmentations (Leung et al. 2011; Frigo et al. 2021; Pérez-Ortega et al. 2021; Weiser et al. 2021). The threshold for ICs to be defined as spatially matching was 0.5, that is 50% or more pixels overlapped between a pair of binary ICs (see Figure 2B). For plotting purposes, ICs at any timescale matching the template map were assigned the same color identity as the maximally matching (highest Jaccard index) IC in the template. We determined spatial matches for each IC in the timescale maps and the template map. All ICs matching template ICs at each timescale were superimposed to yield a cortex-wide map of percentage occurrence for the template ICs in smaller timescales.

The second analysis identified ICs present in the timescale-specific maps that were not present in the template map. Catalogs of timescale-specific ICs were generated by creating a library of all IC binary masks from all ICA segmentations within a timescale. Dimensionality reduction of the IC libraries was achieved with t-distributed stochastic neighbor embedding (t-SNE) using the Jaccard distance algorithm, plotting the position of each IC in this space as a point in a 2D graph (Matlab 2019 *tsne* function). Next, the reduced IC libraries were clustered using Gaussian mixture models, solving for 1 to 100 clusters. The best performing model and cluster number were chosen by minimizing the Bayesian Information Criterion. Occasionally, the best Gaussian mixture model allowed for similar ICs segregating into multiple clusters. ICs from each cluster were superimposed and any IC areas occurring in less than 15% of the time-windows were discarded to achieve an average IC shape and subsequently converted into a binary mask. Remaining ICs containing less than 250 contiguous pixels were also discarded. The catalogs of ICs were matched back to the template ICs using the Jaccard index with a 0.5 match threshold to remove ICs matching the template set. The catalogs of timescale-specific ICs were also matched back to themselves to remove ICs similar to one another, yielding a final map of timescale-specific ICs. Catalogs of timescale-specific ICs were then treated as the new reference map and matched back to each time-window at their timescale using the Jaccard index and threshold as previously described.

### Comparison of ICA segmentations across behaviors

To determine if any ICs are specific to certain behaviors, we concatenated bouts of resting, walking, or grooming that lasted for five seconds or longer and performed ICA for each behavior as described for the timescale-specific datasets.

### Statistical analyses

Statistical analyses were performed using GraphPad Prism 8 (San Diego, CA). The data sets were assessed for normality using a D’Agostino Pearson or Sharpiro-Wilks test (if the n was less than 8). As many data sets were not normally distributed, we used non-parametric statistical tests, including Friedman and Krusal-Wallis ANOVAs. Data are displayed as mean ± standard deviation. Custom Matlab codes for performing pre-processing, ICA, matching of ICs, and behavioral segmentation are available upon request.

## Results

### Database

We recorded mesoscale cortex-wide Ca^2+^ activity from Thy1-GCaMP6f mice (n = 6), while head-fixed on a freely moving disk treadmill that allowed for spontaneous walking, resting, and evoked grooming. The imaging dataset for each mouse was analyzed at several timescales including as a whole and over days, trials, minutes, and 30 second windows. Mice were imaged an average of 565.7 ± 235.7 minutes.

### Functional parcellation of the cerebral cortex based on the entire data set

Initially, for each mouse, all data were concatenated chronologically, and ICA was performed on the combined dataset (Figure 1A). The resulting ICs were used as a ground-truth “template map” to which all other IC segmentations were compared (Figure 1B; Figure S1A). Across mice (Figure 1B; Figure S1A), the cerebral cortex is mostly covered by the 33 ± 9 template ICs. While there are overall similarities in the ICs, the exact template map differs for each mouse, thereby providing an individualized segmentation based on the recorded neuronal dynamics.

**Figure 1:**
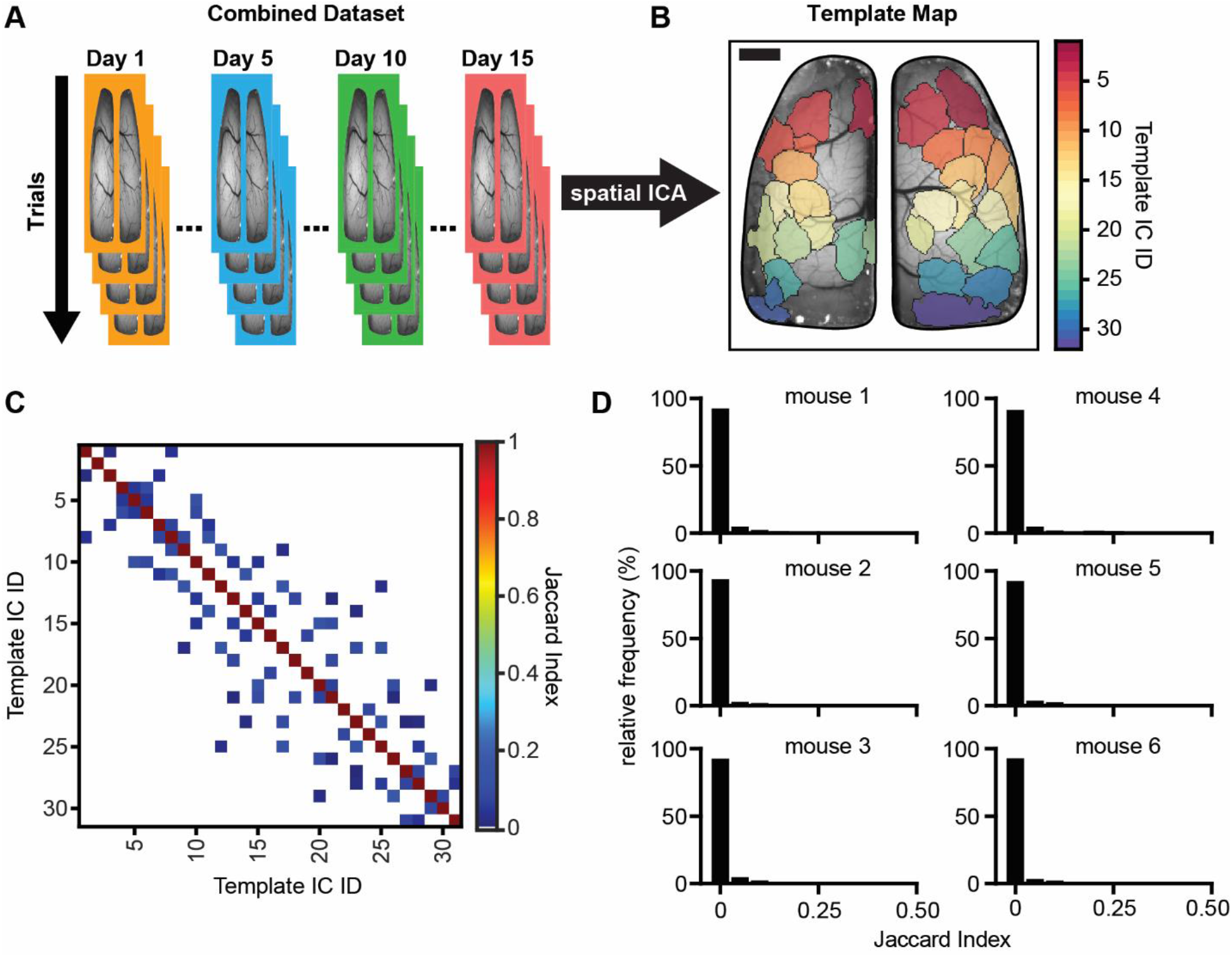
Spatial independent component analysis (ICA) of wide-field calcium imaging data produces spatially independent brain regions. **A)** Schematic showing the ICA workflow of concatenating data for each animal chronologically across days and trials (days are signified by different colored borders; trials are signified by overlapping images) and sending the combined dataset through the JADE ICA algorithm. **B)** Example template map (ground-truth to which all other ICA solutions are compared) of spatial independent components (ICs) produced from running ICA on one mouse’s combined dataset (each different colored region is a single IC; scale bar 1 mm). **C)** Example matrix of Jaccard indices comparing the template map to itself (low off-diagonal Jaccard indices indicate good spatial separation; zero values are shown as white indicating no IC overlap). **D)** Frequency histograms showing the distribution of off-diagonal Jaccard indices (non-self matches) when comparing the template map to itself for each animal (bin-widths = 0.05).

To validate use of the Jaccard index for comparing the similarity of ICs and the matching threshold (≥ 0.5), we computed the Jaccard index for all pairs of template ICs (Figure 1C). Given how ICA segments the data, we expected that the ICs would be maximally spatially independent and IC pairs would have a very low Jaccard index (Rutledge and Bouveresse 2013). The Jaccard index for the vast majority of off-diagonal ICs pairs is 0 (shown as white spaces in an example Jaccard matrix; Figure 1C) and only a small incidence of positive Jaccard indices were observed (blue squares off the diagonal, Figure 1C). The frequency distribution of Jaccard indices across all mice reinforces that there is very little spatial overlap among template ICs, with the vast majority of pairs having no overlap (Figure 1D). Across all mice, the average Jaccard index between template ICs was 0.0063 ± 0.02 (mean ± SD) with an overall maximal value of 0.26, demonstrating that template maps of ICs based on the entire data set are highly spatially separate, therefore confirming that the Jaccard index is a valid measure of IC spatial homology.

### Functional parcellation of the cerebral cortex and comparison on multiple timescales

Using the Jaccard index as our measure of similarity, we investigated the degree to which template spatial ICs were observed over progressively smaller timescales. As described in the Methods, the data were divided into day (trials recorded on a single day), trial (5 or 5.5 minute periods of continuous data), minute (1200 continuous frames), and 30 second (600 continuous frames) periods. Each time segment of data was then run through the ICA algorithm to produce timescale-specific cortical segmentations. The process is shown in Figure 2A, in which all ICs for three individual days are plotted. Next, the Jaccard index was calculated between all pairs of the template set of ICs and the ICs generated for each timescale (Figure 2B). Varying levels of overlap were observed as shown for several examples (Figure 2C), finding that a Jaccard value of 0.5 provides a suitable threshold for a spatial match at the different timescale ICs. Using this threshold, template matching ICs are present at all timescales (Figure 2D), although the number of matches decreases at the smaller timescales.

**Figure 2:**
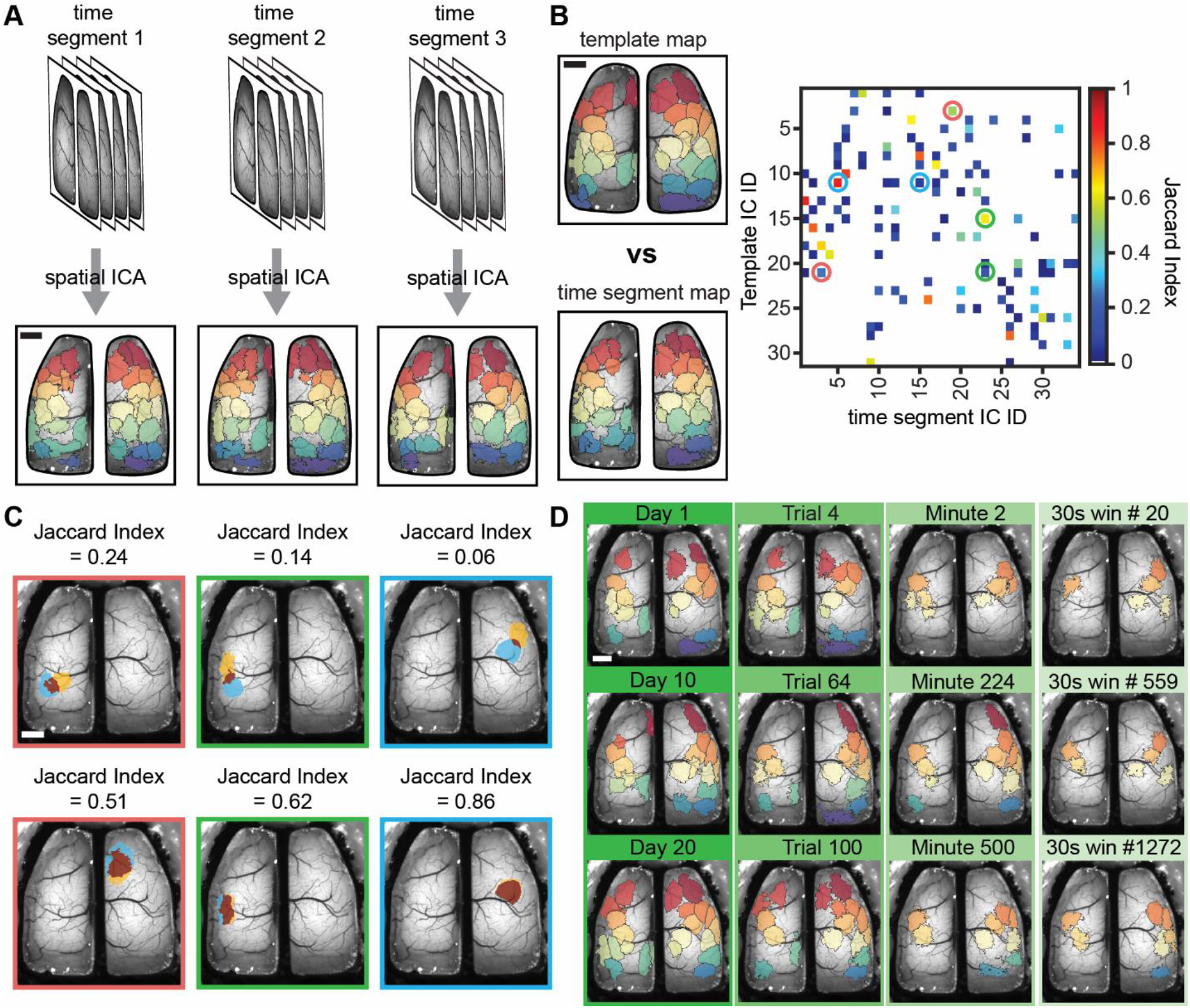
Segmentation of calcium imaging data reveals spatially similar ICs across time segments and scales. **A)** Schematic showing how data is parsed into equal length time segments (set of images show data time segments) for different timescales which are run through the JADE ICA algorithm independently to obtain new ICA solutions (lower images; each color represents an IC). **B)** Example template map based on an entire mouse’s dataset and time segment ICA map at the day timescale (left; colors show individual ICs) and Jaccard index matrix showing matching (high value) and non-matching (low value) ICs between a time segment and template ICA map (circles). **C)** Examples of overlap between matching and non-matching pairs of template and time segment ICs for circles shown in the Jaccard matrix of B (overlap shown in dark red; individual ICs blue or yellow). **D)** Three examples of template matching ICs (Jaccard index ≥ 0.5) for each of the four timescales (columns/green bars). Scale bars: 1mm.

### Template ICs are present at all timescales, but the probability of occurrence varies

Having established a criterion for determining the similarity of ICs across different timescales, we asked how stable the ICs are across time, determining the probability of occurrence of each template IC at the different timescales (Figure 3A). Results reveal that template ICs are timescale invariant, as the same ICs are observed at all timescales. However, the probability of extracting a template IC depends on the duration of the time window analyzed. The average probability of IC occurrence decreased as the duration of time segments was reduced (Figure 3B, % occurrence Days: 55.45 ± 33.19%, Trials: 38.41 ± 34.93%, Minutes: 17.82 ± 26.24%, 30s: 10.19 ± 18.25%; mean ± SD; n = 198; p < 0.0001 Friedman ANOVA; p <0.0001 all post-hoc Dunn’s comparisons). Importantly, the decrease in IC occurrence is not uniform across the timescales or the cortex, demonstrating that the reduction is not simply due to smaller time segments being analyzed. Instead, the occurrence rate is highly dependent on IC location. At the day timescale, many of the ICs are found at 100% probability, and at the minute and 30 second timescales, ICs within sensorimotor regions have occurrence rates of about 80% (Figure 3A and B). Conversely, for ICs in the most anterior and posterior regions, the occurrence probability is less than 20% at the minute and 30 second timescales.

**Figure 3:**
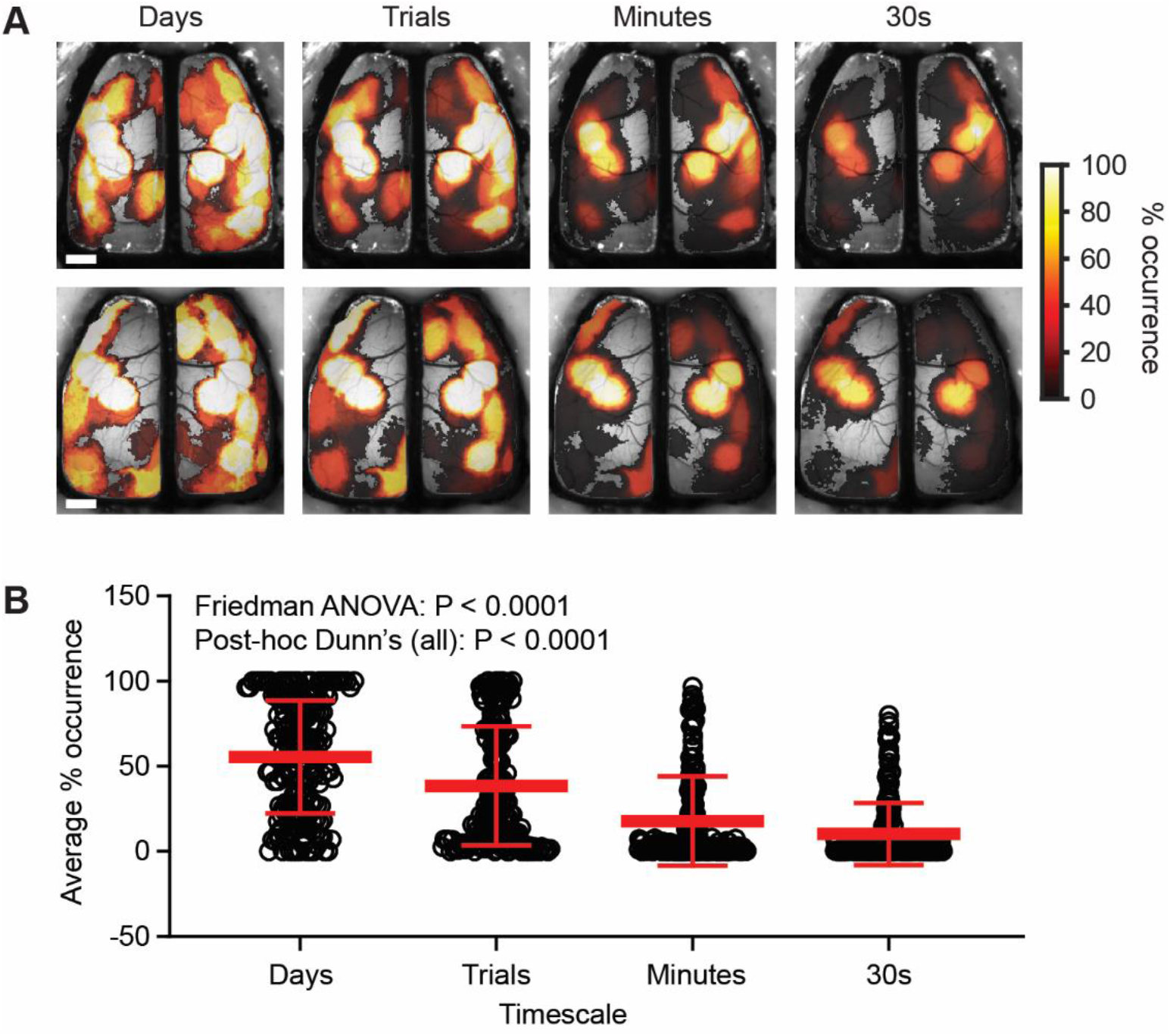
Template ICs are revealed to have a variable temporal presence in brain function as timescales shorten. **A)** Example heat maps from two mice showing the percentage occurrence of template ICs as a function of time at four progressively shorter timescales. **B)** Average percentage occurrence of template ICs across animals at each timescale (Friedman ANOVA p<0.0001; post-hoc Dunn’s comparisons (all) p < 0.0001). Scale bars 1 mm.

The template ICs with the highest probability of observation are in the sensorimotor areas (Figure 3B). Conversely, ICs covering more anterior secondary motor regions and posterior parietal/visual regions are less likely to be present. These results indicate while the same neuronal ensembles that underlie the ICs are present at each timescale; the probability of occurrence is highly dynamic.

To compare the overall cortical coverage at a timescale, we collapsed the template matching ICs at each timescale to create a binary coverage map (Figure 4A). The collapsed maps reveal that the timescale-specific ICs cover a similar area of the cortex as the template ICs. Comparing the binary maps of cortical coverage revealed a high degree of similarity, as evident in the high Jaccard indices between maps at each timescale (Figure 4B; off diagonal comparisons). These findings show that the underlying Ca^2+^ activity can be segmented into a common set of ICs, irrespective of the timescale analyzed.

**Figure 4:**
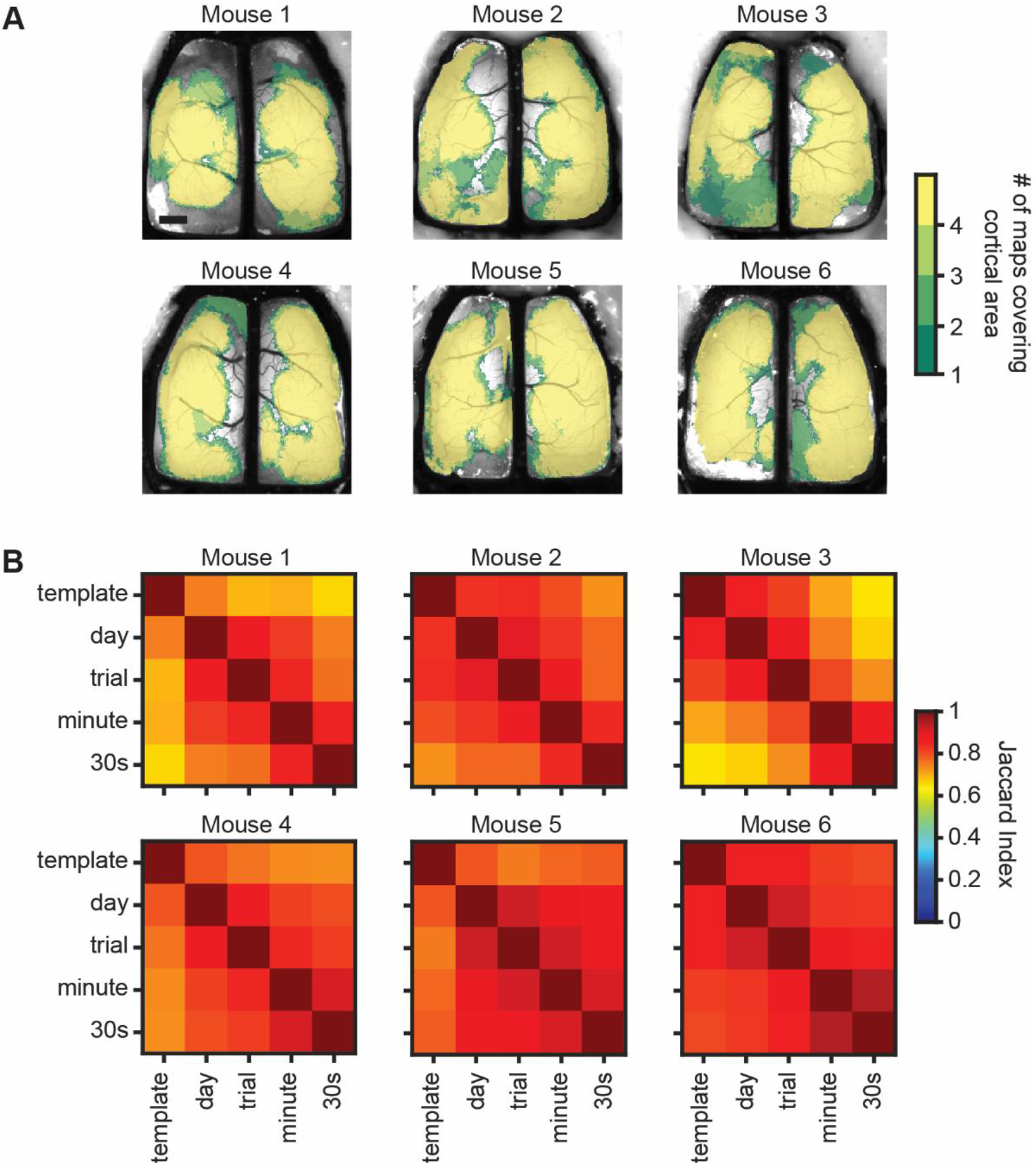
Cortex-wide maps cover similar areas across timescales. **A)** Brain maps from all six experimental subjects showing cumulative cortical coverage of template matching ICs across all time-windows within each of the four timescales examined (color scale shows the number of timescales where an area of cortex was covered by a spatial IC). Scale bar: 1 mm. **B)** Jaccard index matrices for each of the six experimental animals showing a high degree of overlapping cortical coverage between timescales (off-diagonal comparisons).

### Sectioning of data into discrete timescales reveals timescale-specific ICs

The template map for each mouse represents the ICs extracted over a large data set, as typically undertaken with ICA. This raises the question whether new ICs occur at different timescales that are not found in the complete data set. Therefore, we combined all the ICs within a timescale to determine whether unique ICs emerge at shorter timescales. We utilized Gaussian mixture models to cluster ICs within a timescale to identify repeated ICs, as detailed in the Materials and Methods and illustrated in Figure S2. Using the Jaccard index, we then matched the timescale-specific clusters of ICs back to the template set of ICs for each mouse and removed template matches, resulting in timescale-specific ICs.

Intriguingly, timescale-specific ICs were observed across all time windows. These unique ICs occur throughout both hemispheres, with most emerging in anterior motor regions and in parietal and visual cortices (Figure 5A). These same unique ICs can be present across each timescale, as shown in the probability of occurrence plot (Figure 5B). Similar to the template ICs, timescale-specific ICs showed reduced rates of occurrence at the smaller time segments (Days: 34.13 ± 18.97, Trials: 24.05 ± 19.33, Minutes: 14.93 ± 15.10, 30 seconds: 12.33 ± 14.7; Kruskal-Wallis ANOVA p <0.0001; n = 22-75 ICs). These data suggest that unique ICs are relatively common at the day and trial timescales, and show the flexibility of cortical segmentation in both space and time.

**Figure 5:**
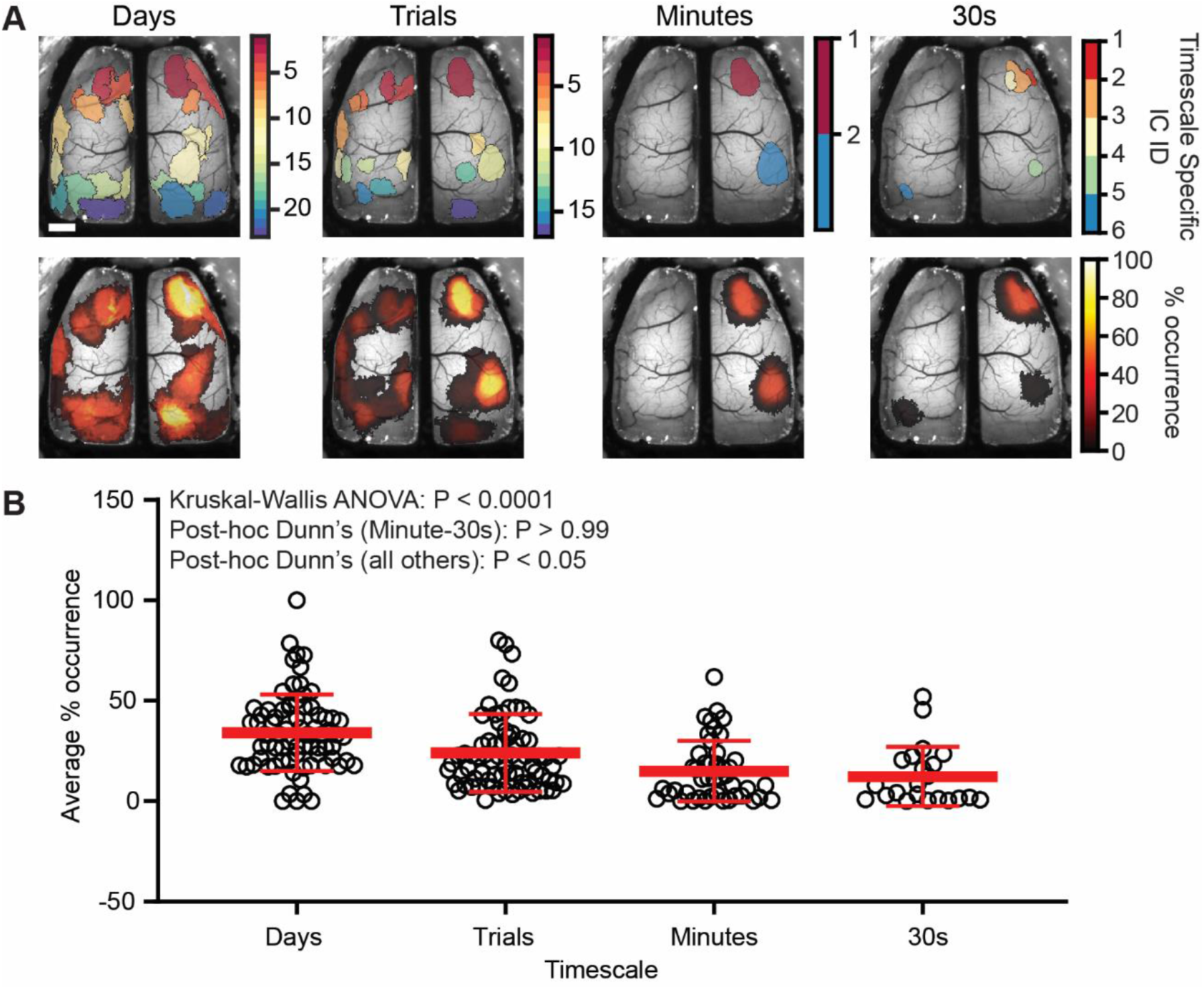
Timescale-specific ICA solutions produce ICs that are unique from the template map. **A)** Example timescale-specific IC maps at each of the four timescales tested (top) and heat maps of the percentage occurrence for timescale-specific ICs for one mouse (bottom). **B)** As the timescale shortens, fewer timescale-specific ICs are produced and are also less frequently observed over time (Kruskal-Wallis ANOVA p < 0.0001; Dunn’s post-hoc comparisons; Minutes-30 s p > 0.99; p < 0.05 all others).

### Different behaviors have both common and unique ICs

The above ICA analyses of the functional segmentation reveal a set of timescale invariant ICs as well as transient, timescale-specific ICs. Therefore, we hypothesized that segmentation based on rest, walk, and grooming would reveal both common and behavior-specific ICs. Behavior-specific maps were generated by performing ICA on datasets that consisted of only rest, walking, or grooming (Figure 6A). ICs in the behavioral maps were matched with those in the template map (see Materials and Methods; Figure 6B right). On average, approximately 60% of the ICs in the template map were represented in the behavior-specific maps (Rest: 64.5 ± 9.5%; Walk: 61.8 ± 13.6%; Groom: 59.6 ± 13.9%; n = 4-6; Figure 6B left). While the matching ICs are located throughout the cortex, each behavior is characterized by a different set of matching ICs. Conversely, there are unique ICs which participate in specific behaviors (Figure 6C). The percentage of behavior-specific ICs was greater for walking (33.06 ± 15.56%) and grooming (32.72 ± 13.93%) than for rest (16.62 ± 6.4%). As with the template matching ICs, behavior-specific ICs occur throughout the cortex and each behavior exhibits a different set of unique ICs. The relatively high percentage of specific ICs, particularly for walking and grooming, highlights the behavior-dependent nature of cortical functional segmentation.

**Figure 6:**
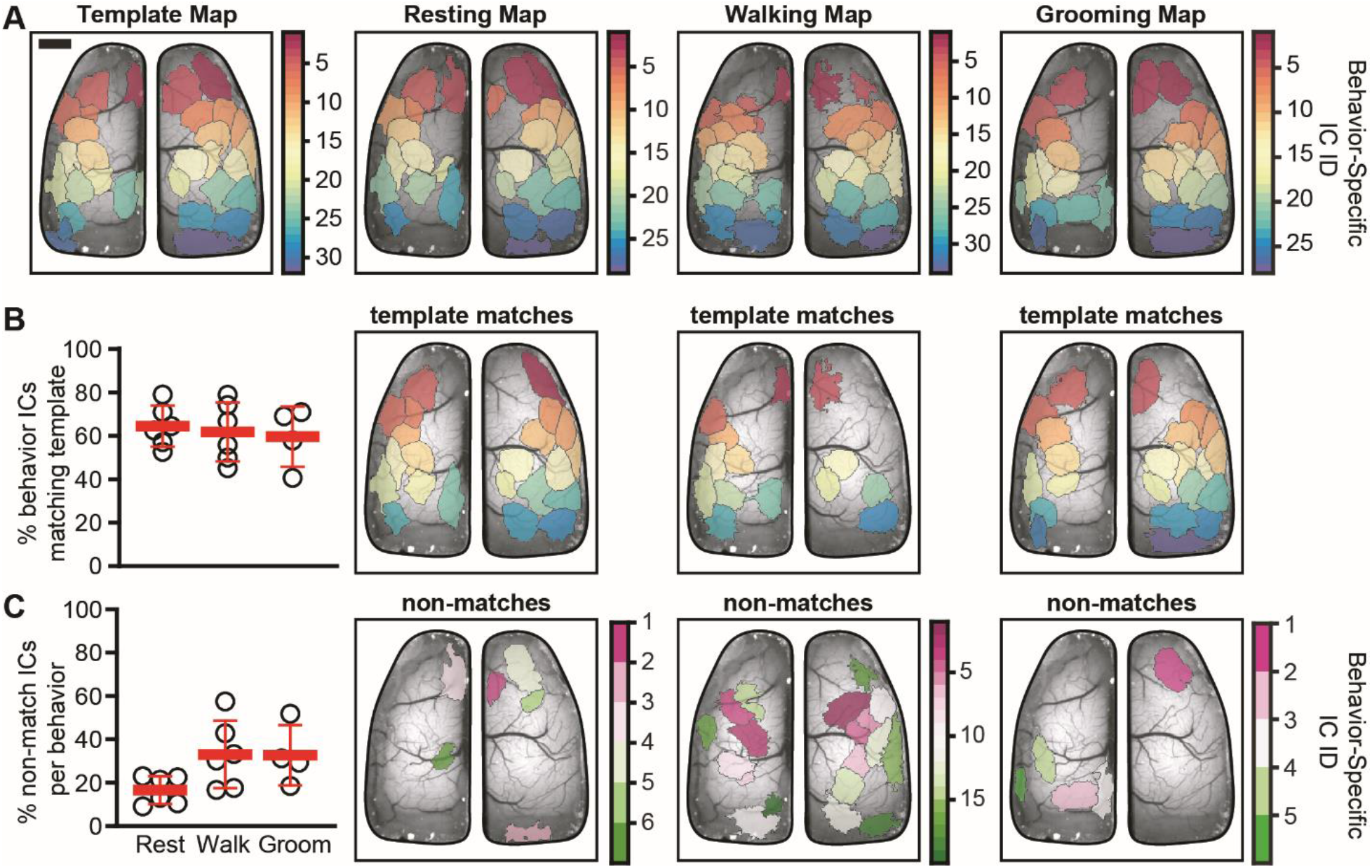
ICA segmentation by behavior produces both template and behavior-specific ICs. **A)** Examples of the template map (left) and behavior-specific IC maps for resting, walking, and grooming. **B)** Scatter plot showing the percentage template ICs that have matches in the behavior-specific IC maps (left) and examples of ICs in each behavior-specific map matching the set of template ICs (right). **C)** Scatter plot showing the percentage of unique ICs generated as part of the behavior-specific maps (left) and examples of behavior-specific ICs generated during each behavior (right). Scale bar: 1 mm.

Finding that both template and behavior-specific ICs participate in cortical processing during behavior, we sought to determine how versatile template ICs are with respect to specific behaviors compared to the full dataset irrespective of behavior. To do this, Jaccard indices were calculated between all pairs of ICs in the template map and each behavior map. If a template IC had a matching IC in the behavior-specific map, it suggests that the IC participates in the that behavior. ICs in the template map were then color coded according to which behavior or combination of behaviors in which they were engaged (Figure 7). Results show that a majority of template ICs are engaged in multiple behaviors with 39.2 ± 10.1% in all three, 7.2 ± 3.6% in rest and walk, 12.1 ± 11.5% in groom and rest, and 5.9 ± 4.1% in groom and walk. Fewer template ICs are specific to a single behavior with 12.4 ± 10.5% in rest, 13.8 ± 10.9% in walk, and 6.9 ± 3.2% in groom. Lastly, 13.2 ± 5.5% of template ICs are not engaged during any of the behaviors analyzed, suggesting these ICs are active during different stimuli and/or behaviors. The high rate of IC engagement across behaviors implies that a core set of regions are frequently used to encode behavior.

**Figure 7:**
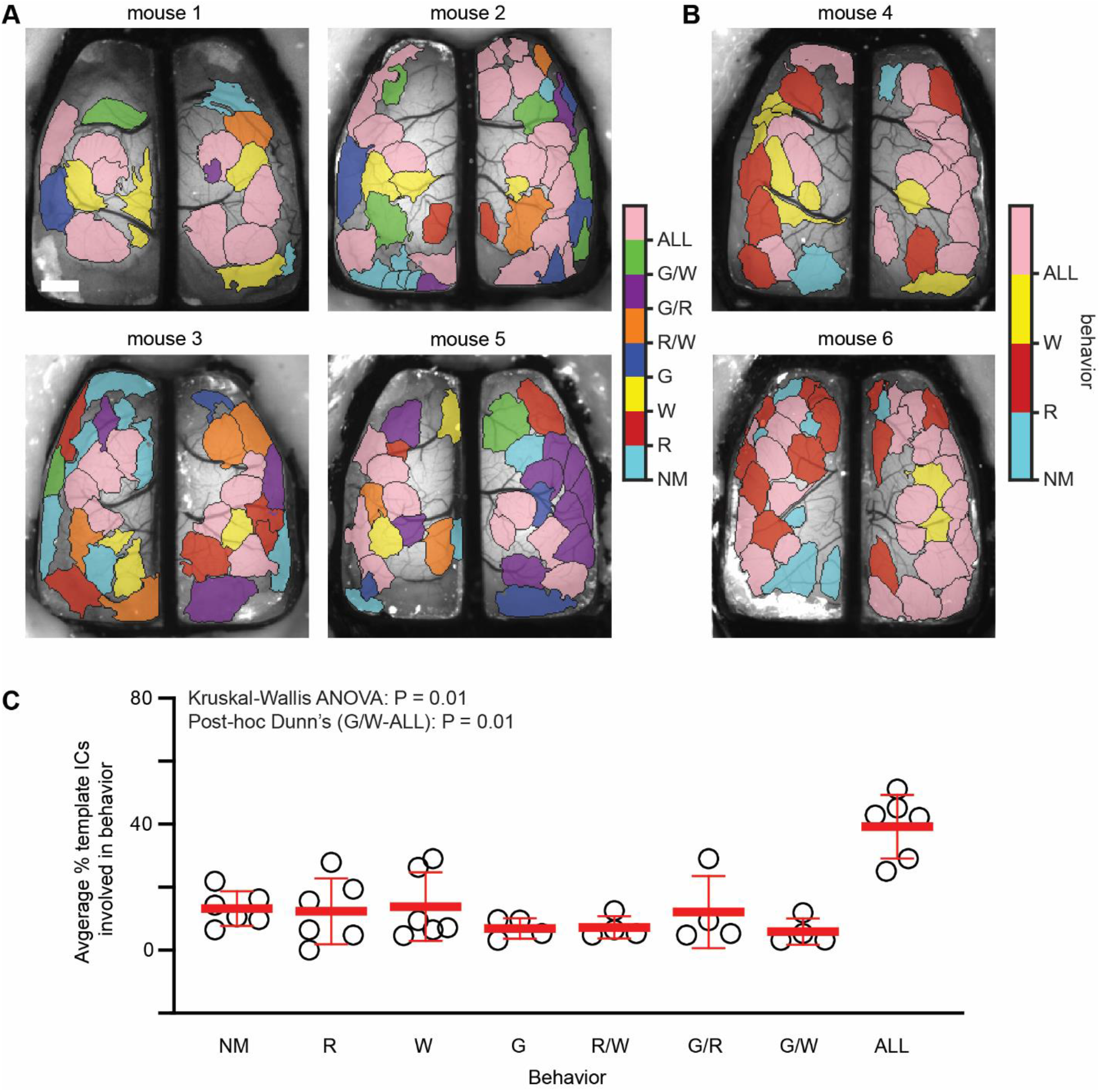
Template map ICs are engaged in a behaviorally specific manner. **A)** Template IC maps for four animals with all three behaviors showing the behavior or combination of behaviors in which each IC participates. Each color corresponds to a specific behavior or combination of behaviors. **B)** Similar to A, template IC maps for two animals in which grooming behavior was not present. **C)** Scatter plot showing the percentage of template ICs involved in each behavior or combination of behaviors. The only significant difference in % ICs was ALL versus G/W (Kruskal-Wallis ANOVA p = 0.01; post-hoc Dunn’s comparisons G/W-ALL p = 0.01, all others p > 0.05). NM - No match, R - Rest, W - Walk, G - Groom, R/W - Rest and Walk, G/R - Groom and Rest, G/W - Groom and Walk, ALL - All behaviors. Scale bar: 1 mm.

Finally, we investigated which behavior-specific map ICs are shared across behaviors. As with previous results, we hypothesized that ICs would be both shared between and unique to specific behaviors. When comparing the resting behavior map to both walking and grooming behavior maps, we find that approximately 50% of ICs are shared between behaviors, although different combinations of ICs are observed in the individual behaviors (Figure 8A). Similarly, when comparing the walking or grooming specific maps to the other behaviors, we find both shared and unique ICs; which changed depending on the behavior observed (Figure 8B-C). As shown in for the example mouse in Figure 8A-Cii, comparing the segmentation between behaviors reveals a set of shared ICs as well as unique ICs. A similar finding was observed across all mice tested (Figure 8A-Ciii). The shared ICs between behaviors range from 44.4 to 58.5% and unique ICs with a single behavior range from 35.2 to 55.6% (Table 1). These data suggest that a core map of ICs participates in generating all behaviors and is reconfigured on an as needed basis. These data also suggest that while many of the same regions are engaged across behaviors, new cortical areas are integrated in a behavior-specific manner, increasing computational flexibility.

**Figure 8:**
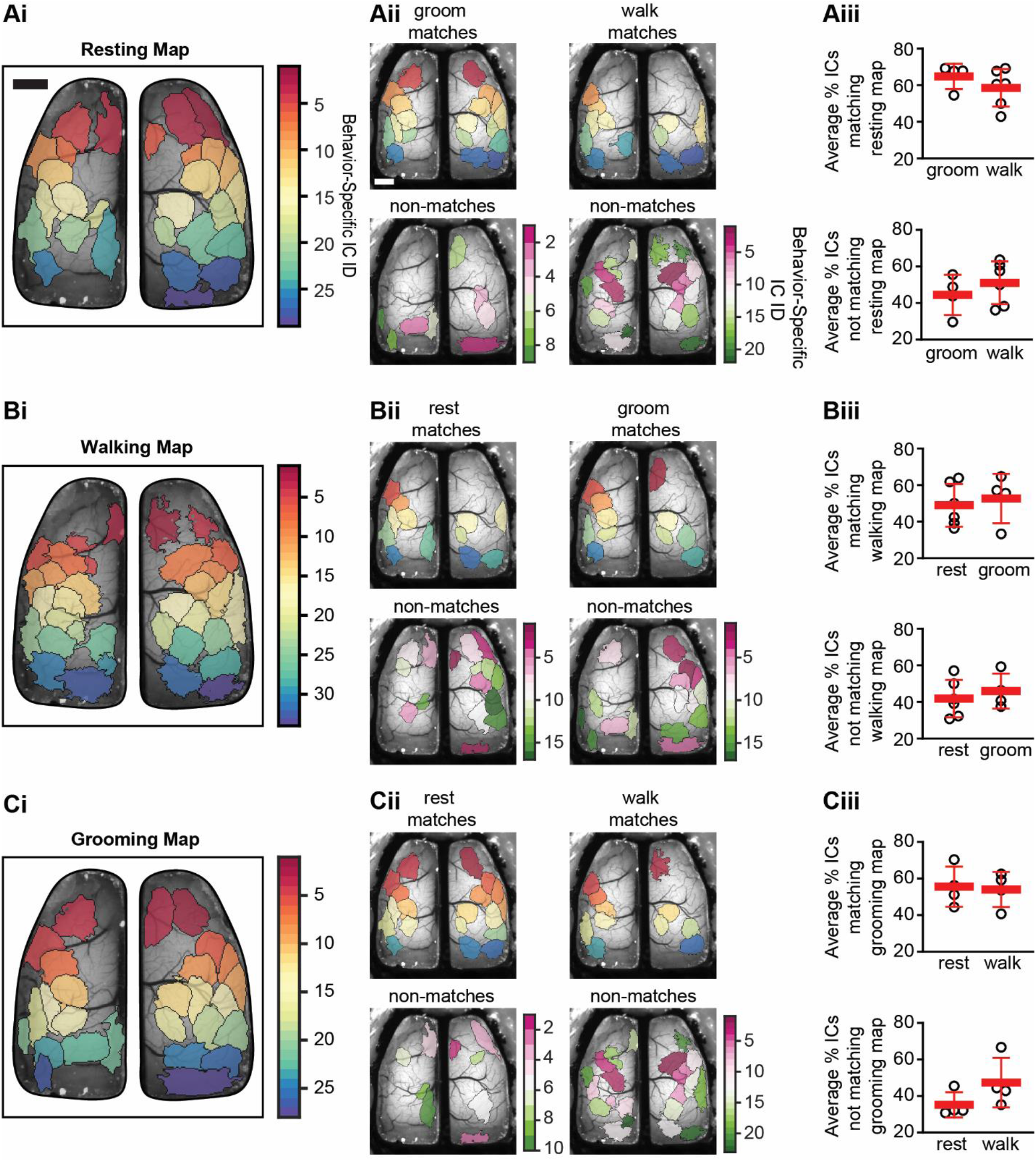
Behavior-specific ICA segmentations generate both shared and unique ICs. **Ai-Ci)** Example spatial ICA domains obtained by combining data from resting, walking, or grooming behaviors which is used as reference map for comparing between behaviors. **Aii-Cii)** Example maps of matching (top) and unique (bottom) ICs when comparing all behaviors. **Aiii-Ciii)** Scatter plots showing the percentage of ICs matching between behaviors (top) and percentage of ICs unique from the reference behavior (bottom). Scale bars: 1 mm.

**Table 1:** Summary of shared and behavior-specific ICs across subjects. Using behavior-specific maps as the reference map shows approximately half of ICs are shared across behaviors whereas approximately half are behavior-specific. Data shown are mean ± SD. Percentage shared and unique ICs for each behavior map were compared using Mann-Whitney (MW) tests. In all MW tests p>0.05.

| Reference Map | Rest ICs Shared (%) | Walk ICs Shared (%) | Groom ICs Shared (%) | Rest Specific ICs (%) | Walk Specific ICs (%) | Groom Specific ICs (%) |
| --- | --- | --- | --- | --- | --- | --- |
| Rest | N/A | 58.5 ± 10.3 | 64.8 ± 6.9 | N/A | 51.1 ± 11.7 | 44.4 ± 11 |
| Walk | 48.6 ± 11.7 | N/A | 52.7 ± 13.5 | 41.8 ± 10.2 | N/A | 46 ± 9.6 |
| Groom | 55.6 ± 11 | 54 ± 9.6 | N/A | 35.2 ± 6.9 | 47.3 ± 13.5 | N/A |

## Discussion

To understand the neural processes measured with cortex-wide mesoscale Ca^2+^ imaging we used spatial ICA, a blind source method that segments the imaged area into functional regions with minimal overlap based on statistically independent neuronal Ca^2+^ activity. To gain insight into the dynamics and relevance of the segmentation, we partitioned the data into a wide range of timescales and discrete behaviors, including rest, walk, and grooming.

The key findings are that a common set of ICs are stable, being present across a wide range of timescales, ranging from 30 seconds to days. In addition to the common ICs in the template, unique ICs emerge at the shorter duration timescales. Segmenting the data by behavior reveals both common and behavior-specific ICs, with unique combinations characterizing rest, spontaneous locomotion, and grooming. Together, these findings suggest that the functional segmentation within the cerebral cortex is spatially stable, but also that neural assembles possess the ability to form and dissolve as needed, for example, during specific behaviors.

### ICs are stable across timescales

The observation that the template ICs are present at the shorter timescales and cover the same area of the dorsal cerebral cortex has several implications. First, the template ICs represent functional regions of the cortex that are consistently engaged in these head-fixed, behaving mice. This stability of the template ICs over timescales suggests that different behaviors use the same functional groups of neurons (see below).

Second, the probability maps of the template ICs obtained across the day to 30 second timescales may be analogous to the default mode and behavioral networks observed with human functional imaging (Kiviniemi et al. 2011; Lu et al. 2012; Stafford et al. 2014). The ICs with the highest probabilities across timescales are the primary motor and somatosensory cortices, followed by retrosplenial and secondary motor area, all regions which would be expected to be engaged during the behaviors observed. The highest occurrence rates for ICs in the somatomotor regions are consistent with the observation that ongoing cortical activity is driven by the diverse array of ongoing spontaneous motor activity in head-fixed mice (Gilad et al. 2018; Musall et al. 2019; Salkoff et al. 2020; Ren and Komiyama 2021).

Conversely, we postulate that the template ICs with lower occurrence rates represent cortical regions involved in other behaviors, variations on the more common behaviors, responses to sensory stimuli, or changes in brain states. As shown previously, head-fixed mice engage in a large repertoire of movements and behaviors that modulate neural activity that would account for time-dependent engagement of template ICs (Drew et al. 2019; Musall et al. 2019; Salkoff et al. 2020).

### Timescale-specific ICs reveal flexibility in cortical segmentation

While we show that ICA performed over large data sets yields a stable functional segmentation, we also observed timescale-specific ICs that were relatively common at the day, trial, and even minute timescales. Timescale-specific ICs are of interest, as they are transient and reflect subtle but important changes in the spatial organization of activity, which are masked when analyzing longer time periods. This finding mirrors similar results on simulated and human EEG and fMRI data using a sliding window ICA approach, showing that ICA source maps become more dissimilar as the timespan between segmentations increases (Jung et al. 2001; Esposito et al. 2003; Kiviniemi et al. 2011). These studies, in addition to the present results, show that cortical segmentation is flexible over time. With recent interest in inter-individual IC map variability and its effect on functional connectivity mapping, the dynamic maps generated at smaller timescales may prove crucial for accurate activity mapping (Bijsterbosch et al. 2019; Bergmann et al. 2020).

Intriguingly, the timescale-specific ICs were located in the secondary motor areas and in posterior parietal and visual processing areas. These more anterior and posterior placed ICs tend to surround the somatomotor regions defined by the template ICs that emerge most frequently. One possibility is that these transiently activated regions reconfigure the core network for specific processes. The anterior motor areas covered by timescale-specific ICs are involved in planning and decision making (Svoboda and Li 2018; Inagaki et al. 2022; Xu et al. 2022; Yin et al. 2022). The visual areas and posterior parietal regions are critical for both vision and sensory-motor integration (Beloozerova and Sirota 2003; Freedman and Ibos 2018; Steinmetz et al. 2019; Wal et al. 2021). These processes are likely subject to changes in the behavioral context, as opposed to the somatomotor regions that are covered with high probability by the template ICs and are involved in more stereotypical aspects of behaviors (Xu et al. 2022).

### Behavior-specific maps are a combination of shared and unique ICs

In the initial analysis, we investigated the temporal dynamics of the functional modules in a behaviorally agnostic manner. The observation of timescale-specific ICs suggests that cerebral cortical segmentation dynamically engages different neuronal ensembles. To test this concept, we partitioned the data into three specific behaviors: rest, walk, and groom. We observed that a majority of the template ICs are present in all three behaviors. However, the same template ICs are not found in each behavior, instead each behavior engages a subset of the template ICs. Further, during rest, walk and groom, new ICs are extracted. Therefore, each behavior is characterized by a different functional segmentation. The relatively high percentage of behavior-specific ICs, particularly for walking and grooming, highlights the dynamic nature of cortical segmentation that would not be captured in a static cortical map. The implications are that the cerebral cortex uses both common and unique regions to encode and execute the full repertoire of behaviors.

### ICA provides robust functional segmentation

As outlined in the Introduction, the development of wide field imaging techniques represents a major advance for characterizing cortex-wide interactions during behavior but comes with the major challenge of massively high dimensional datasets. Saxena and colleagues recently outlined several desiderata for the analysis of wide-field imaging (Saxena et al. 2020). These include: 1) denoising and compression, 2) avoiding non-physiological constraints such as the orthogonality required by SVD, 3) decomposition into interpretable signals defined by well-defined regions, 4) flexibility to adapt to differences across animals, 5) is comparable across animals, and 6) reproducibility. Overall, we demonstrate that ICA meets, if not exceeds, each of these criteria for functional segmentation of wide-field Ca^2+^ imaging data.

The resulting ICA yielded a template map with on average 33 ICs across mice. These ICs were obtained from large data sets of mesoscale Ca^2+^ imaging data after denoising and compression. As a blind source separation technique, ICA decomposes a complex, multi-channel signal into a linear set of independent components, with the goal of identifying the underlying sources and their time courses. ICA has many compelling features and had been suggested to be a “more principled decomposition of biological time series” (Friston 1998). Depending on the implementation, the components can be either temporal or spatial. As used in this study, spatial ICA finds the spatial sources of the Ca^2+^ fluorescence in the cerebral cortex based on high order statistics and non-Gaussian features of the data. While similar to principal component analysis (PCA) or singular value decomposition (SVD), the components obtained using PCA or SVD are orthogonal. The orthogonality of the linear bases derived from PCA and SVD has always been questioned as problematic, if not unrealistic, in biological systems. However, in ICA the components are both orthogonal and uncorrelated. Therefore, ICA is a strong approach to identify the underlying sources in the data and avoids non-physiological constraints such as parcellation based on orthogonality.

As ICA is a data-driven approach without any assumptions about the spatial structure, ICA can adapt to most differences across animals. This would include both local or generalized lesions, pathology, developmental differences, and experimental localizations. In each animal, a unique set of ICs are determined, based on the underlying neuronal activity. The ICs are well-defined regions, typically single cortical regions. When an IC is composed of two non-contiguous regions, they are most often a pair of homotopic cortical areas. This is consistent with the activity in the two areas being tightly coupled via the corpus callosum. Furthermore, ICs can be compared across animals.

Assessing the reproducibility across timescales and behaviors was one of the main foci of the present study. We found remarkable reproducibility as highly spatially similar ICs were extracted over timescales ranging from hours to 30 second subsections and across recording sessions that spanned, on average, 19 days. While clearly meeting the desired reproducibility, the spatial stability observed implies that the ICs operate as functional units over both shorter and longer duration time scales. When all of these properties are taken together, spatial ICA provides a robust cortical segmentation to investigate the network structure under a variety of conditions.

### Limitations of ICA

Several authors have pointed out some of the limitations of ICA. Although we, and others, have emphasized that the functional segmentation derived from ICA is data-driven, one concern was that the approach is not hypothesis driven or well-suited for statistical modeling (Friston 1998). However, the ICs can certainly be used to test hypotheses of interest (McKeown and Sejnowski 1998).

Each IC has a value across the entire space and, therefore, a threshold must be set to define the spatial boundaries (Sui et al. 2009; West et al. 2021). This typically is done on the Z-score of the ICs. Here we used a z-score of ± 2.5 SD and the Jaccard index shows this yields spatial ICs with almost no overlap. Using a conservative threshold may be problematic, yielding a segmentation into components that does not capture the functional integration among brain regions (Friston 1998). Relaxing the Z-score threshold, would produce larger ICs, with more spatial overlap. However, as we observed previously using JADE, the spatial profile of cortical ICs are characterized by either one or two, very steep maxima (West et al. 2021). A widely accepted approach to setting the threshold remains to be determined.

The spatial ICA segmentation is unique for each animal and the ICs vary in location across the cortex, as would be expected for an activity-based methodology. This poses challenges to averaging results across animals. Analysis across animals is easier when using a fixed segmentation. In human imaging, this has been addressed by using group ICA methods like temporal concatenation group-ICA and probabilistic ICA. However, the use of these group-wise techniques has been limited for wide-field Ca^2+^ imaging (Calhoun et al. 2001; Hui et al. 2011; Cramer et al. 2019). As many ICA algorithms are agnostic to data-type, group-ICA of wide-field Ca^2+^ imaging data is a potential solution to comparing ICs across mice.

### Implications for cerebral cortical processing and network dynamics

As we show using ICA for wide-field Ca^2+^ imaging, the cerebral cortex has the capacity for both stable and dynamic segmentation. This finding carries major implications for network analytics like functional connectivity (FC), as generating functional networks on large datasets (e.g. using template ICs) would not capture either the temporal or behavior-specific connectivity. Furthermore, changing functional segmentation in time and with behavior may reveal unique functional networks engaging in cortical processing, as temporal variability of template and unique ICs suggests that the network is reconfigured on an as-needed basis. The ability to selectively utilize brain regions may allow for more efficient information processing.

The finding that template ICs vary with time and that new segmentations emerge and dissolve with different behaviors allows for a larger dynamic range of neural network possibilities than what would be dictated by anatomy alone. Furthermore, this spatial diversification of information coding would make cortical networks both stable and resistant to failure by creating redundant pathways for processing (Huber et al. 2012; Clopath et al. 2017). This temporal and structural variability shows similarity to computational neural networks that are most optimal when operating on a semi-stable basis (Rössert et al. 2015; Hochstetter et al. 2021). Similar properties have also been observed at the single cell level in neural ensembles (Huber et al. 2012; Lütcke et al. 2013; Pérez-Ortega et al. 2021). Together, these findings suggest that this property of microscale flexibility, coupled with macroscale stability, exists across multiple levels of processing. This dynamic stability allows the brain to distinguish environmental or behavioral differences while still retaining information as learned behaviors or contexts. Our timescale and behavior-specific ICA analyses reveal that while the cortex possesses a remarkable stability in functional organization, it is also capable of a high level of flexibility in this functional organization, depending on the task at hand.

## Acknowledgements

We would like to thank Lijuan Zhuo for her assistance with the animal surgeries. The Minnesota Supercomputing Institute provided high-processing computing, and the University of Minnesota University Imaging Centers (UIC, SCR_020997) provided 3D printing services of the cortical implants. This work was supported in part by National Institutes of Health grants, R01 NS111028 and P30 DA048742 to T.J.E., and K99 NS121274 to M.L.S., as well as by an AIRP grant from the University of Minnesota Medical School to T.J.E.

