## Supplementary figures for "To be and not to be: Wide-field Ca^2+^ imaging reveals neocortical functional segmentation combines stability and flexibility"

### Supplementary Material

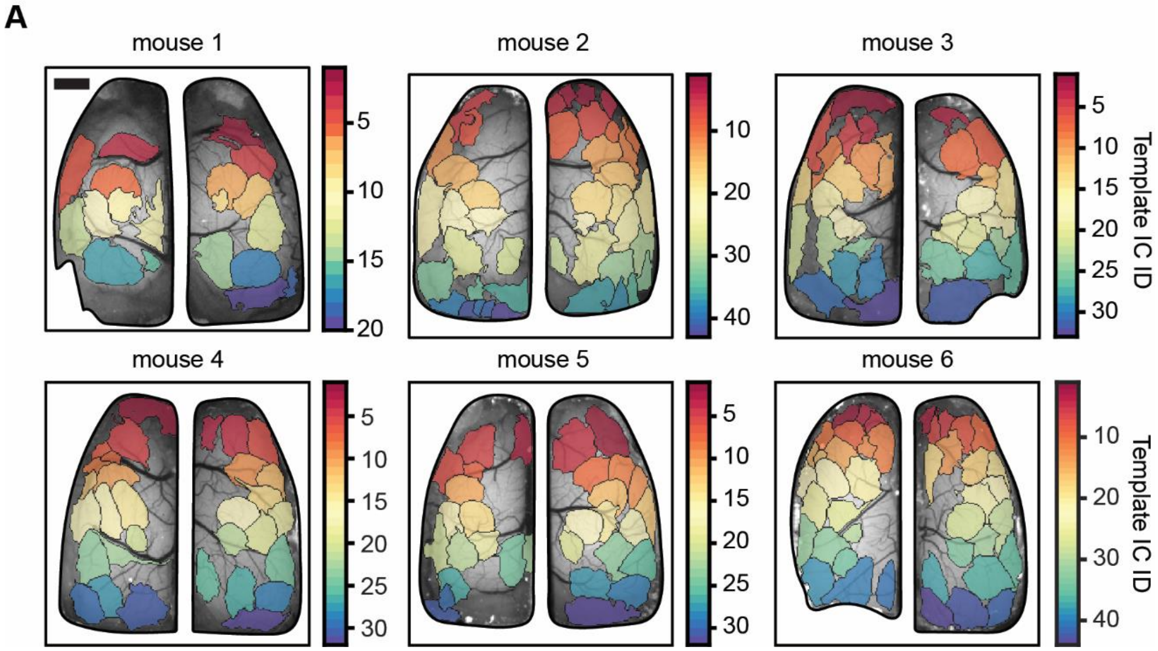


**Supplementary Figure 1: Spatial ICA produces animal specific template maps. A)** Template IC maps for each animal included in this study after running spatial ICA on the combined dataset for each mouse. Each colored region is an independent region; artifacts not shown. Scale bar: 1 mm.

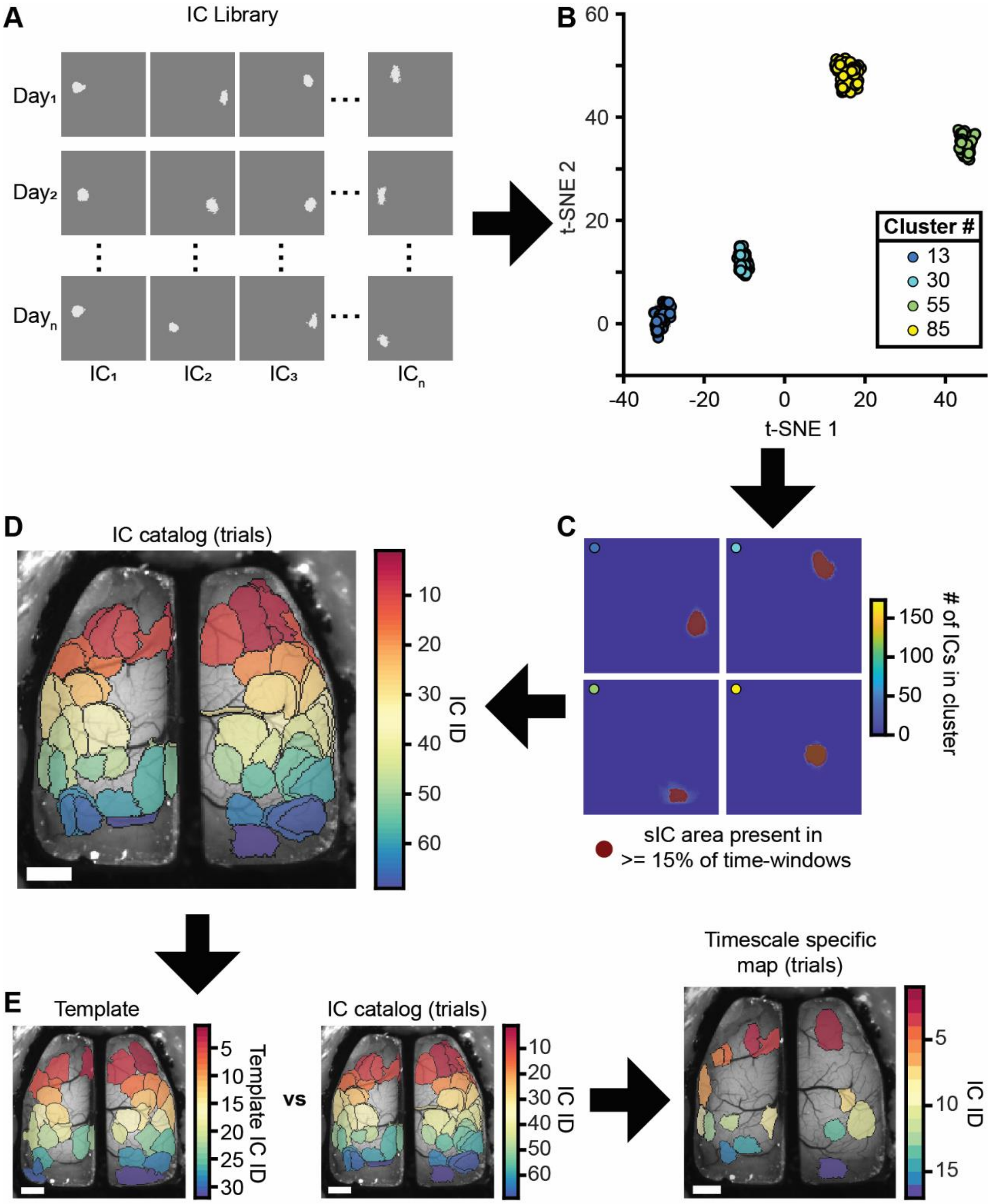


**Supplementary Figure 2: Timescale-specific ICA solutions reveal sets of unique spatial ICs. A)** Timescale-specific libraries of ICs are created by combining all spatial ICs for each time window of a timescale. **B)** Timescale-specific spatial IC libraries are reduced to 2 dimensions and ICs similar in spatial location and shape are clustered together using Gaussian mixture models. **C)** Clustered spatial ICs are superimposed and an average shape of the clustered ICs is obtained by removing regions appearing in less than 15% of time windows analyzed for each timescale. **D)** Spatial ICs remaining after step C are converted to binary images and yield a timescale-specific catalog of ICs. **E)** The timescale-specific catalog of spatial ICs is matched to the template (to remove template ICs) and also back to itself (to remove spatially similar ICs that separated into multiple clusters). The remaining spatial ICs make up the timescale-specific IC map which is composed of ICs unique to analysis at that timescale (right). Scale bars: 1 mm.
